# Biallelic Variants in *MAD2L1BP* (*p31^comet^*) Cause Female Infertility Characterized by Oocyte Maturation Arrest

**DOI:** 10.1101/2023.01.16.524217

**Authors:** Lingli Huang, Wenqing Li, Xingxing Dai, Shuai Zhao, Bo Xu, Fengsong Wang, Ren-Tao Jin, Lihua Luo, Liming Wu, Xue Jiang, Yu Cheng, Jiaqi Zou, Caoling Xu, Xianhong Tong, Heng-yu Fan, Han Zhao, Jianqiang Bao

**Affiliations:** Reproductive and Genetic Hospital, The First Affiliated Hospital of USTC, Division of Life Sciences and Medicine, University of Science and Technology of China, Hefei, Anhui, 230001, China; Hefei National Laboratory for Physical Sciences at Microscale, Biomedical Sciences and Health Laboratory of Anhui Province, University of Science and Technology of China (USTC), Anhui, China; Life Sciences Institute, Zhejiang University, Hangzhou 310058, China; Center for Reproductive Medicine, Cheeloo College of Medicine, National Research Center for Assisted Reproductive Technology and Reproductive Genetics, Key laboratory of Reproductive Endocrinology of Ministry of Education, Key laboratory of Reproductive Endocrinology of Ministry of Education, Shandong University, Shandong, China; School of Life Science, Anhui Medical University, Hefei 230022, China

**Keywords:** MAD2L1BP, oocyte maturation arrest, meiosis, spindle assembly checkpoint (SAC), infertility

## Abstract

Human oocyte maturation arrest represents one of the severe conditions for female patients with primary infertility. However, the genetic factors underlying this human disease remain largely unknown. The spindle assembly checkpoint (SAC) is an intricate surveillance mechanism that ensures accurate segregation of chromosomes throughout cell cycles. Once the kinetochores of chromosomes are correctly attached to bipolar spindles and the SAC is satisfied, the MAD2L1BP, best known as *p31*^*comet*^, binds MAD2 and recruits the AAA+-ATPase TRIP13 to disassemble the mitotic checkpoint complex (MCC), leading to the cell cycle progression. In this study, by whole-exome sequencing (WES), we identified homozygous and compound heterozygous *MAD2L1BP* variants in three families with female patients diagnosed with primary infertility owing to oocyte metaphase I (MI) arrest. Functional studies revealed that the protein variants resulting from the C-terminal truncation of MAD2L1BP lost their binding ability to MAD2. cRNA microinjection of full-length or truncated MAD2L1BP uncovered their discordant roles in driving the extrusion of polar body 1 (PB1) in mouse oocytes. Furthermore, the patient’s oocytes carrying the mutated *MAD2L1BP* variants resumed polar body extrusion (PBE) when rescued by microinjection of full-length *MAD2L1BP* cRNAs. Together, our studies identified and characterized novel biallelic variants in *MAD2L1BP* responsible for human oocyte maturation arrest at MI, and thus prompted new therapeutic avenues for curing female primary infertility.

## Introduction

Infertility is increasingly becoming a global health issue that affects ∼10% to 15% (e.g., approximately 186 million) of couples at reproductive age around the world (Inhorn and Patrizio, 2015). Currently, many couples benefit from assisted reproductive techniques (ARTs), such as *in vitro fertilization* (IVF) or *intracytoplasmic sperm injection* (ICSI), to have their own babies(Beall et al., 2010a). However, among the patients seeking ART treatment in the clinic, some of the females suffer from severe recurrent failure even through repeated IVF/ICSI attempts. These female patients with primary infertility were most often attributed to functional deficiency in the oocytes, such as oocyte maturation arrest, premature oocyte death, fertilization failure and preimplantation embryonic arrest(Beall et al., 2010b; Mehlmann, 2005). Oocyte maturation is referred to as meiotic resumption of oocytes from the germinal vesicle (GV) to the metaphase II (MII) stage *in vivo* or *in vitro*, whereby the oocytes acquire the developmental competence. In the clinic, oocyte maturation arrest is characterized by the frequent occurrence of oocytes mostly or entirely arrested at the germinal vesicle (GV) or metaphase I (MI) stage, representing a rare but highly challenging disease for treatment through traditional ART(Beall et al., 2010b; Mehlmann, 2005; Mrazek and Fulka, 2003).

Unlike the male germline, female germ cells initiate meiosis very early during embryonic development once primordial germ cells (PGCs) transit and colonize the genital ridge in mice and humans. Nonetheless, following the synapsis and recombination between homologous chromosomes, meiotic progression ceases in the embryonic gonad, and the oocytes remain arrested at the diplotene stage in meiotic prophase I during postnatal oocyte growth in growing follicles. After birth, a limited number of primordial follicles are activated and recruited from the primordial follicle pool to develop sequentially into primary follicles, secondary follicles, and antral follicles. This process is concurrent with the oocyte cytoplasmic and meiotic maturation process whereby the oocytes continuously grow with increased size and acquire the ability to resume meiotic division upon luteinizing hormone (LH) surge (Maddirevula et al., 2017b; Mrazek and Fulka, 2003). The fully grown oocyte is morphologically characterized by the appearance of germinal vesicles, representative of the oocyte nucleus with a bulgy nucleolus that can be readily identified under phase-contrast microscopy. Morphologically, meiotic resumption is characterized by chromatin condensation and the dissolution of nuclear membranes, termed “germinal vesicle breakdown” (GVBD). This is followed by the cell-cycle progression to MI and subsequent extrusion of the first Polar Body (PB1), culminating in metaphase II (MII) arrest, which is ready for fertilization(Lane and Kauppi, 2019; Mihajlovic and FitzHarris, 2018). This prolonged meiotic prophase I arrest at diplotene stage in conjunction with the discontinuous, hormone-triggered progression to MII stage in oocytes is highly genetically regulated and spatiotemporally coordinated. In agreement with this conclusion, numerous genetically engineered mouse models (GEMMs) have uncovered the genetic factors that are required to drive accurate and timely meiotic progression. For instance, genetic ablation of *Cdc25b* or *Pde3a* led to GV arrest, while deletion of *Mei1, Cks2*, or *Mlh1* caused MI arrest in mouse oocytes(Mehlmann, 2005; Mihajlovic and FitzHarris, 2018; Mrazek and Fulka, 2003; Nguyen et al., 2018; Touati et al., 2015; Wassmann et al., 2003). Occasionally, genetic abrogation can cause a mixed phenotype. For example, mouse oocytes deficient in *Mlh3* arrested at both the MI and MII stages, whereas *Ubb-null* oocytes ceased at both the GV and MI stages (Lenzi et al., 2005; Lipkin et al., 2002). Although these mouse genetic models provide insightful mechanisms underlying oocyte meiosis, the causative deleterious variants responsible for oocyte maturation arrest resulting in human infertility remain scarce. Until recently, only a limited number of protein-coding variants have been discovered to account for human oocyte maturation arrest, including *TUBB8*(Feng et al., 2016) (MIM: 616768), *PATL2*(Chen et al., 2017; Huang et al., 2018; Maddirevula et al., 2017a) (MIM:614661), and *TRIP13*(Zhang et al., 2020) (MIM: 604507). The malfunction of these genes often caused the persistent activation of the oocyte spindle assembly checkpoint (SAC) owing to improper disassembly of mitotic checkpoint complex (MCC), resulting in oocyte maturation arrest. However, mutations in these genes can only account for a very small proportion (<30%) of patients with this syndrome, and the deleterious causative variants in most cases remain largely unknown(Chen et al., 2017; Christou-Kent et al., 2018; Feng et al., 2016; Huang et al., 2018; Maddirevula et al., 2017a; Zhang et al., 2020). Therefore, there is an urgent need to screen the causal factors among the female patients suffering from recurrent reproductive failure for better genetic diagnosis and treatment in the future.

## Results

### Identification of biallelic variants in *MAD2L1BP* underlying human oocyte maturation arrest at the MI stage

A cohort of 50 female patients with primary infertility displaying the oocyte maturation arrest at the MI stage was recruited from the Reproductive Center of the First Affiliated Hospital of USTC (University of Science and Technology of China) and the Reproductive hospital at Shandong University between July 2014 and October 2021. All the patients had normal karyotypes (46, XX) and had undergone at least one cycle of IVF or ICSI treatment, IVF or ICSI treatments of the three patients with variants in *MAD2L1BP* are shown (Table 1). The studies were approved by the ethics committees of the University of Science and Technology of China and Shandong University. The oocytes and blood samples were donated by individuals who had provided the written, informed consent paperwork. The DNA samples were subjected to conventional WES screening in order to underpin the genetic variants. We screened the candidate variants based on the following criteria: (a) rare variants with a minor allele frequency (MAF) below 1% in five public databases: 1000 genome, dbSNP, gnomAD, EVS, and Exome Aggregation Consortium (ExAC); (b) nonsynonymous exonic or splice site variants, or frameshift INDEL; (c) heterozygous variant that is also carried by the parents; (d) known RNA expression in our *in-house* oocyte expression database); (e) homozygous variants were prioritized in consanguineous families (Figure 1A, see also Materials and Methods). This pipeline analyses led us to narrow down the candidate variants to a short list in three families (12, 14 and 11 candidate genes in the respective families) (Table 2, Table 2-source data 1), and eventually to focus on one shared gene variant, namely *MAD2L1BP*, wherein two homozygous nonsense variants (c.853C>T [p.R285*] and c.541C>T [p.R181*]) of *MAD2L1BP* from two unrelated families (Family 1 and Family 3), respectively, and two compound heterozygous variants (a frameshift mutation p.F173Sfs4* and an intronic mutation c.21-94G>A) of MAD2L1BP in Family 2 (Table 2-source data 1, Table 3).

**Table 1.** Clinical Characteristics of Affected Individuals and Their Retrieved Oocytes.

| Age<br>(Years) | Duration of<br>Infertility (Years) | IVF/ICSI<br>Cycles | Protocol | Total Number of<br>Oocytes Retrieved | GV<br>Oocytes | MI<br>Oocytes | PB1<br>Oocytes | Fertilized<br>Oocytes | Cleaved<br>Embryos |
| --- | --- | --- | --- | --- | --- | --- | --- | --- | --- |
| Family 1 (II-1) |  |  |  |  |  |  |  |  |  |
| 22 | 2 | 1(IVF) | Modified ultra-long | 11 | 0 | 11 | 0 | 0 | 0 |
|  |  | 2(IVF) | Mild stimulation | 6 | 1 | 5 | 0 | 0 | 0 |
|  |  | 3(IVF) | PPOS | 13 | 0 | 13 | 0 | 0 | 0 |
| Family 2 (II-1) |  |  |  |  |  |  |  |  |  |
| 31 | 6 | 1(IVF) | PPOS | 9 | 0 | 9 | 0 | 0 | 0 |
| Family 3 (II-1) |  |  |  |  |  |  |  |  |  |
| 34 | 9 | 1(IVF) | Natural | 1 | 0 | 1 | 0 | 0 | 0 |
|  |  | 2(IVF) | Short | 4 | 0 | 4 | 0 | 0 | 0 |
| GV, germinal vesicle; MI, metaphase I; PB1, first polar body. PPOS, Progestin-primed ovarian stimulation. |  |  |  |  |  |  |  |  |  |

**Table 2.** Summary of the variants identified after filtering by WES in patients from three families (F1: II-1, F2: II-1 and F3: II-1)

|  | F1: II-1 | F2: II-1 | F3: II-1 |
| --- | --- | --- | --- |
| Total variants | 134159 | 392392 | 66914 |
| After excluding variants reported in dbSNP, 1000 genomes, EVS, ExAC and gnomAD. (MAF < 0.01) | 3619 | 14023 | 22832 |
| Exonic non-synonymous or splicing variants, or coding indels | 273 | 571 | 9660 |
| Homozygous or compound heterozygous (excluding X and Y chromosomes) | 12 | 14 | 11 |
| Homozygous | 6 | 5 | 3 |
| In homozygous region > 2Mb | 1 | - | - |

**Table 3.** MAD2L1BP Pathogenic Variants Observed in the three Families.

| **Families** | **Genomic Position** | **cDNA Change** | **Protein Change** | **Mutation Type** | **SIFT<sup>a</sup>** | **PPH2<sup>a</sup>** | **Mutation Taster<sup>a</sup>** | **NNsplice<sup>a</sup>** | **ASSP<sup>a</sup>** | **gnomAD<sup>b</sup> allele** | **gnomAD<sup>b</sup> homozygotes** |
| --- | --- | --- | --- | --- | --- | --- | --- | --- | --- | --- | --- |
| **1** | Chr6: 43608202 | c.853C>T | p.R285* | Nonsense | NA | NA | D | NA | NA | 3/248598 | 0/248598 |
| **2** | Chr6: 43607867 | c.518delT | p.F173Sfs4* | Frameshift deletion | NA | NA | D | NA | NA | Not found | Not found |
|  | Chr6: 43600837 | c.21-94G>A | - | Splicing | NA | NA | NA | 0.13>0.0<sup>c</sup> | 7.1>2.2<sup>c</sup> | 111/31250 | 2/31250 |
| **3** | Chr6: 43607890 | c.541C>T | p.R181* | Nonsense | NA | NA | D | NA | NA | 2/250850 | 0/250850 |

Abbreviations are as follows: D, Deleterious; NA, Not available. <sup>a</sup> Mutation assessment by SIFT, Polyphen-2 (PPH2), Mutation Taster, NNsplice and ASSP. <sup>b</sup> Frequency of corresponding mutations in all population of gnomAD. <sup>c</sup> The variant c.21-94G>A is predicted to introduce an alternative splice acceptor by NNsplice (the increase of the acceptor site score from 0 up to 0.13) and by ASSP (the increase of the acceptor site score from <2.2 up to 7.1).

**Figure 1.**
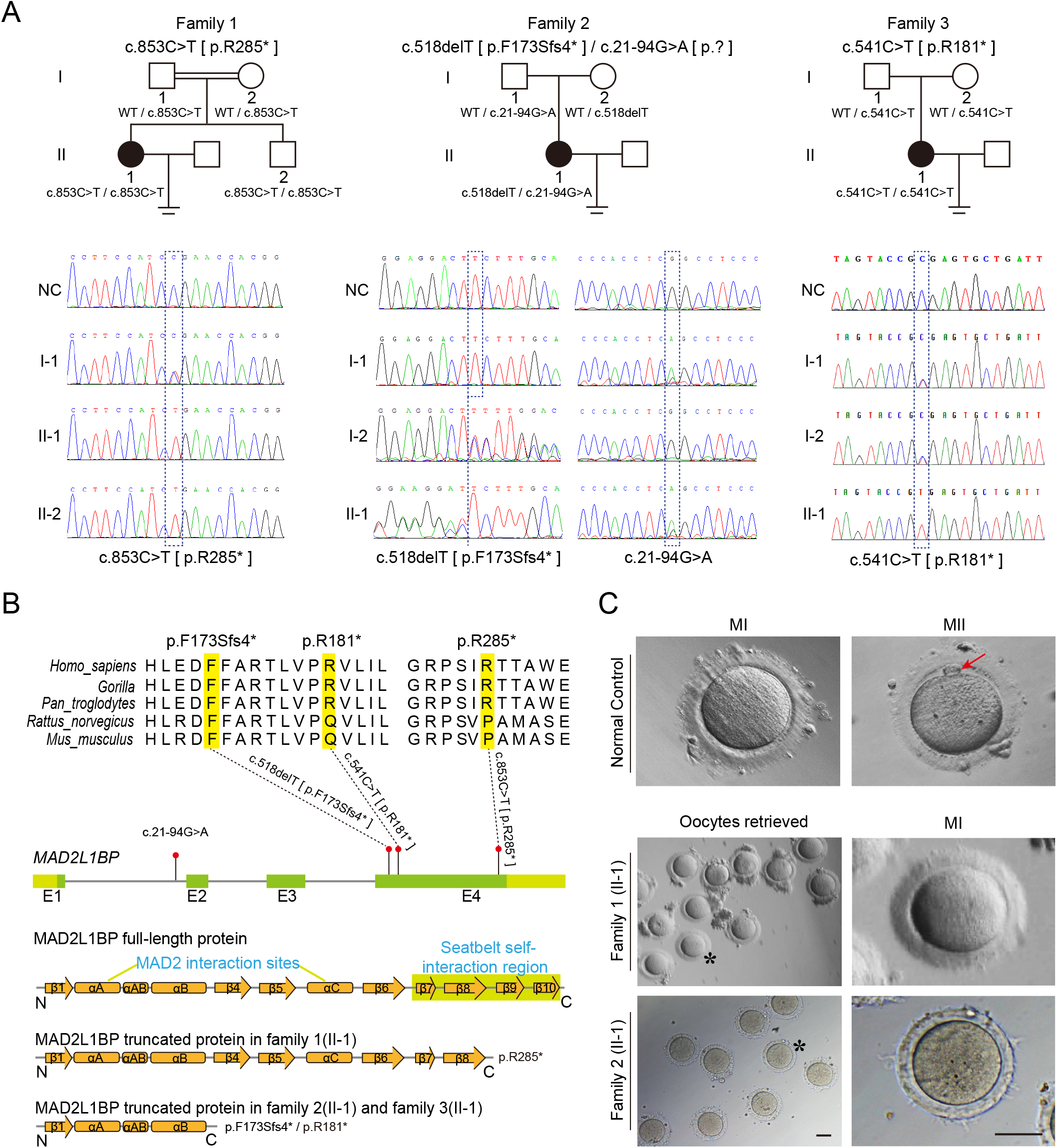
Identification of Pathogenic Variants in *MAD2L1BP* from three unrelated families. (A) Pedigrees of three female patients with oocyte maturation arrest at the MI stage from three unrelated families. The first case was from a consanguineous family, while the other two were not. The patients carried homozygous c.853C>T [p.R285*] and c.541C>T [p.R181*] variants from family 1 and family 3, respectively, while the patient from family 2 had a compound heterozygous mutation c.518delT; c.21-94G>A as indicated. The chromatograms below show the Sanger sequencing result of the PCR-amplified fragments containing the respective variants in each family. (B) Phylogenetic conservation of the identified amino acid mutations in MAD2L1BP. The positions of all mutations are indicated in the schematic genomic structure of *MAD2L1BP*. The MAD2-interacting site and the seatbelt self-interaction region for MAD2L1BP are labelled in the secondary structure of MAD2L1BP protein at the bottom. All identified mutations residing in Exon 4 of *MAD2L1BP* generated the premature STOP codon, likely resulting in truncated proteins. (C) The representative morphology of normal and affected individual oocytes. The normal MI and MII oocytes are shown in the top panel. The red arrow points to the first polar body (PB1). Oocytes retrieved from family 1 (II-1) and family 2 (II-1) were arrested at the MI stage. The asterisks indicate the oocytes with a magnified view in the right panel. MI: metaphase I; MII: metaphase II. Scale bar, 80 μm.

In the first case, the female patient (F1: II-1) was from a consanguineous family and underwent three unsuccessful cycles of IVF with 29 out of a total of 30 superovulated oocytes arrested at the MI stage (Figure 1A, Figure 1C, Table 1). WES analysis following a set of bioinformatic pipelines identified a homozygous nonsense mutation in NM_001003690 (*MAD2L1BP*): c.853C>T [p.R285*] (Figure 1A and 1B). Her younger brother (F1: II-2), 22 years of age, carried the same homozygous mutation (Figure 1A, Figure 1-figure supplement 2). Further Sanger sequencing after PCR amplification confirmed that their consanguineous parents were both heterozygous for this mutation. The second patient was diagnosed with primary infertility with no history of consanguinity. She had gone through one cycle of IVF, and all nine collected oocytes were arrested at MI. (F2: II-1) (Figure 1C, Table 1). A heterozygous frameshift mutation c.518delT [p.F173Sfs4*] in *MAD2L1BP* was uncovered by WES (Figure 1A,Table 3), which was inherited from her mother. Subsequent Sanger sequencing of ∼200 bp intron-flanking sequences of *MAD2L1BP* validated a heterozygous intronic variant c.21-94G>A in the patient (F2: II-1), which was transmitted from her father (Figure 1A-C, Table 3). In the third family, the female patient had a total of five superovulated oocytes all arrested at MI after two unsuccessful cycles of IVF. WES and subsequent PCR sequencing revealed a homozygous mutation c.541C>T [p.R181*] in *MAD2L1BP* (Figure 1A and 1B, Table 1). This mutation was inherited from her parents respectively and likely resulted in the production of a much shorter C-terminally truncated MAD2L1BP protein compared with p.R285* variant from the first patient.

The nonsense variants p.R285* and p.R181* occurred at low allele frequencies (3/248598 and 2/250850, respectively) in the gnomAD browser, and the frameshift mutation p.F173Sfs4* was not found in the database (Table 3). However, the intronic variant c.21-94G>A (rs142226267) had an allele frequency of 111/31250 and homozygote frequency of 2/31250 in the gnomAD browser, which was initially filtered out through the WES analysis pipeline but subsequently reconsidered for further evaluation of its pathogenicity (Table 3). The nonsense variants p.R285* and p.R181* and the frameshift mutation p.F173Sfs4* gave rise to a premature termination codon (PTC) in the resultant *MAD2L1BP* mRNA. For the intronic variant c.21-94G>A, it was predicted to be a cryptic splicing acceptor site of exon 2 by NNsplice and ASSP, presumably leading to aberrant splicing of Exon 2 with additional intronic sequences resulting in a PTC introduction (Table 3).

### C-terminal truncation of MAD2L1BP abolished its interaction with MAD2, a key kinetochore protein

MAD2L1BP is a relatively small-sized protein consisting of only 306 amino acids, which are highly evolutionarily conserved across metazoan species (Figure 1-figure supplement 1 and Figure 2-figure supplement 1). Interestingly, there is a shorter isoform that differs exclusively at the N-terminus compared to the longer isoform in humans (Figure 1-figure supplement 1). By analyzing the published RNA-seq databases, we found that *MAD2L1BP* mRNAs are highly abundant in both human and mouse oocytes, especially those at advanced stages (Figure 2-figure supplement 1), indicative of an important role of MAD2L1BP in oocyte meiotic maturation in mammals. To determine whether those variants impact *MAD2L1BP* mRNA expression, we next designed quantitative PCR (qPCR) primers against *MAD2L1BP* mRNAs. We discovered that p.R285* mutation did not affect the expression levels of *MAD2L1BP* mRNAs in the first patient (Figure 2A), thus possibly producing a truncated MAD2L1BP protein without C-terminal 22 amino acids (Figure 1B). In the second patient carrying the compound heterozygous variants, the p.F173Sfs4* mutation would elicit frame-shift reading causing nonsense-mediated mRNA decay or generate a much shorter C-terminally truncated MAD2L1BP protein. To investigate whether the c.21-94G>A (rs142226267) variant affected splicing of MAD2L1BP mRNA, we initially tried the cDNA amplification using total RNA extracted from the patient’s peripheral blood sample. However, we failed to amplify the predicted bands after several rounds of attempts by optimization of the primer designs and PCR conditions, presumably owing to the extremely low abundance of aberrant MAD2L1BP mRNA in the blood (*data not shown*). Therefore, we next conducted an *in vitro* splicing assay by synthesizing a plasmid that comprised the Exon 1, Exon 2 and Exon 3 as well as the flanking intronic sequences in the *MAD2L1BP* gene. Not surprisingly, the aberrant splicing around Exon 2 was repeatedly observed by sequencing the transcribed spliced mRNA product after transfection into 293T cells *in vitro* (Figure 2-figure supplement 2). In support of this finding, the qPCR assay validated that the *MAD2L1BP* mRNA levels were significantly reduced in the blood from the second patient (Figure 2B). We were unable to quantitatively measure the effect of p.R181* mutation on *MAD2L1BP* mRNA expression in the third patient owing to the unavailability of blood samples. However, the p.R181* mutation would only be possible to generate a much shorter MAD2L1BP protein than p.R285* variant in the first patient, leading to a complete loss of 126 amino acids at the C-terminus (Figure 1B). As such, we chose to use the longer variant (p.R285*) for subsequent functional studies.

**Figure 2.**
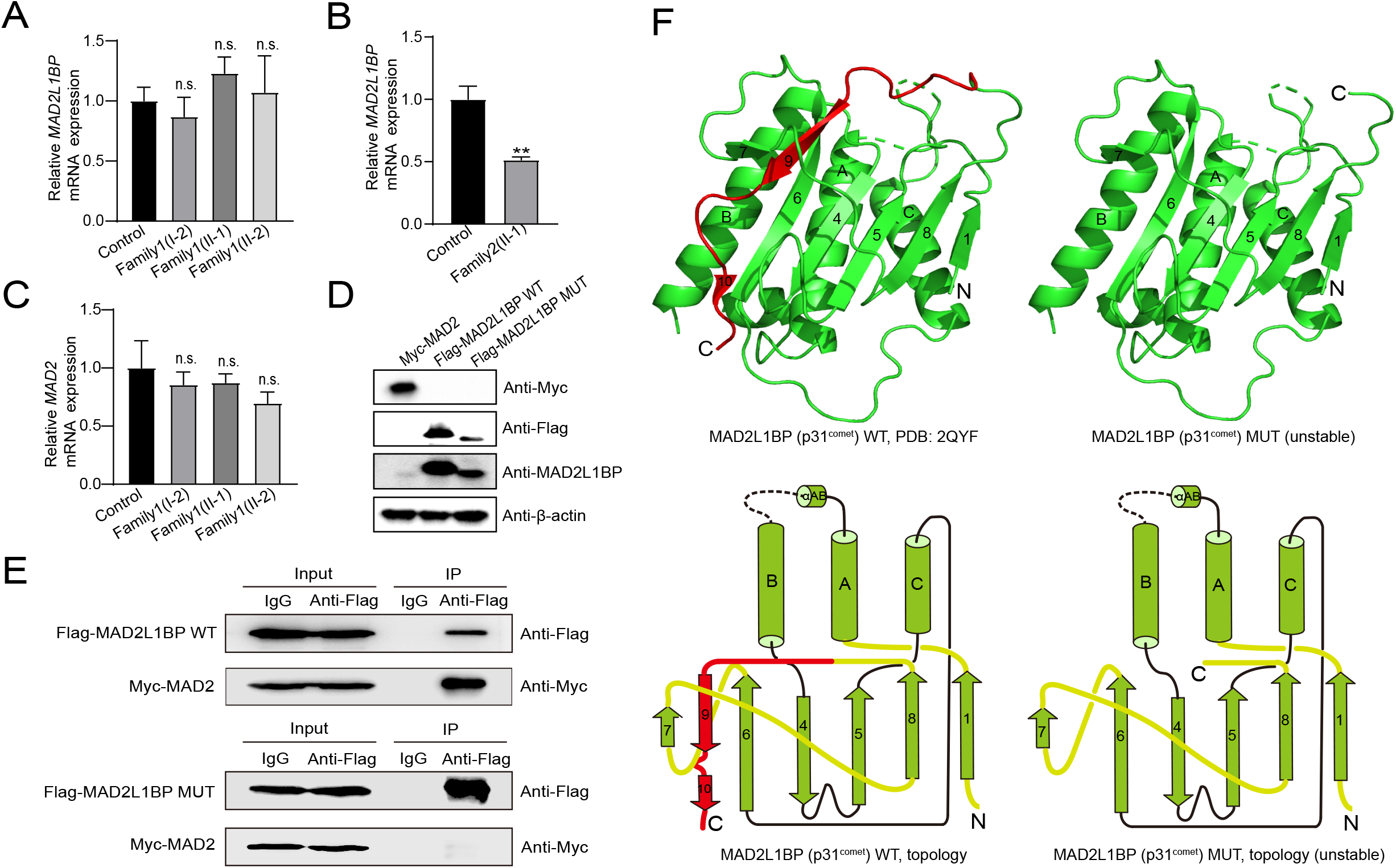
Adverse impacts of *MAD2L1BP* variants on mRNA Expression and Protein function. (A-B) qPCR assays comparing the mRNA expression levels of *MAD2L1BP* in peripheral blood samples from family 1 and family 2 as indicated. The experiments were performed in technical triplicates. (C) qPCR assay showing the mRNA expression levels of *MAD2* in individual blood samples from family 1. The experiments were performed in technical triplicates. (D) The ectopic protein expression levels of Myc-MAD2, Flag-MAD2L1BP WT and Flag-MAD2L1BP MUT (p.R285*) by immunoblotting following transfection of individual plasmids into 293T cells in culture. Full-length and truncated MAD2L1BP protein can be seen in the panel. (E) Co-immunoprecipitation (Co-IP) assays demonstrating that the truncated MAD2L1BP MUT (p.R285*) lost interaction with MAD2, as compared with MAD2L1BP WT, when co-transfected into 293T cells *in vitro*. (F) Ribbon and topological diagrams showing the structure of MAD2L1BP WT (PDB ID: 2QYF) and the predicted MAD2L1BP MUT (p.R285*). Notably, MAD2L1BP MUT (p.R285*) lacks a C-terminal seatbelt configuration (highlighted in red) resulting in structural instability.

Next, we asked whether the C-terminally 22 aa-deleted MAD2L1BP variant owing to p.R285* mutation retained its full function. MAD2L1BP has been shown to act as an adaptor that tightly binds mitosis arrest deficient 2 (MAD2) and recruits TRIP13 (Alfieri et al., 2018; Brulotte et al., 2017; Ma and Poon, 2016). At the protein level, MAD2L1BP harbors highly conserved central “α-helix”- and flanked “β-sheet”-organized domains, which structurally mimick MAD2 and physically interact with each other *in vivo* at the MAD2 dimerization interface (Yang et al., 2007) (Figure 1B and 2F). In accordance with *MAD2L1BP*, the mRNA expression levels of *MAD2* were also high in both human and mouse oocytes (Figure 2-figure supplement 1). MAD2 is a core member of the MCC, which is timely assembled and disassembled through the conformational transition of “O-MAD2” to “C-MAD2” via a “MAD2 template” model throughout the cell cycle (De Antoni et al., 2005). The disassembly of the MCC machinery is predominantly achieved through MAD2L1BP, which acts as an adaptor that binds MAD2 and recruits ATPase TRIP13 to the MCC (Alfieri et al., 2018; Ma and Poon, 2016; Wang et al., 2014). Compelling studies have shown that MAD2 overexpression or deficient TRIP13 rendered oocyte meiotic arrest in both mice and humans, suggesting an indispensable role of MAD2-MAD2L1BP-TRIP13 signaling in driving meiotic division (Wassmann et al., 2003). We therefore subsequently carried out co-immunoprecipitation (Co-IP) assays using Myc-tagged MAD2 plasmid, and Flag-tagged WT and p.R285* mutated MAD2L1BP construct (Figure 1B and 2D). Not surprisingly, while the Flag-tagged WT MAD2L1BP could strongly pull down the full-length MAD2 as judged by Co-IP, the loss of 22 aa in MAD2L1BP at the C-terminus completely abolished its binding to MAD2, suggesting an essential function of the C-terminal 22 aa in stabilizing the binding of MAD2L1BP to MAD2 (Figure 2E). In support of this finding, the resolved crystal structure of MAD2L1BP (PDB: 2QYF) showed that its C-terminal 22 residues fold into the β9 and β10 strands, which are in charge of the physical interaction between MAD2L1BP and MAD2 like a seatbelt essential for the conformational stability (Yang et al., 2007) (Figure 1B and 2F). This evidence altogether suggests that the C-terminus of MAD2L1BP is required for its interaction with MAD2, and the oocyte maturation arrest in the female patients was attributed to the truncated MAD2L1BP variants identified.

### Overexpression of WT Mad2l1bp, but not Mut Mad2l1bp, rescued the mouse oocyte MI arrest *in vitro*

To further corroborate the causal relationship between MAD2L1BP variants and female infertility resulting from oocyte MI arrest, we next interrogated the phenotypic outcome by overexpression of WT or *Mad2l1bp* mutant (hereafter all “mut” represents “p.R285*”) via cRNA injection into PMSG-primed mouse oocytes at the GV stage (Figure 3A). As shown in Figure 3B and 3C, exogenous expression of either WT or p.R285* *Mad2l1bp* alone resulted in similar percentages of oocytes undergoing GVBD. However, further examination in detail showed that the oocytes supplemented with WT *Mad2l1bp* extruded the polar body much earlier than those supplemented with p.R285* *Mad2l1bp* injection. A quantitative comparison from the time-lapse imaging experiments revealed that microinjection of the WT *Mad2l1bp* cRNAs accelerated the polar body extrusion by ∼3 hr as compared with the control group (Non-injected), although the resultant percentages of oocytes with PBE were comparable between both groups (∼80%) (Figure 3B and 3D). By comparison, supplementation with p.R285* *Mad2l1bp* cRNAs not only postponed the extrusion of the polar body but also significantly lowered the final PBE rate in the oocytes (∼40%) (Figure 3B and 3D). This evidence strengthened that the mutant p.R285* is a detrimental mutation rendering *loss of function* for the full-length Mad2l1bp. Subsequently, we eliminated endogenous Mad2l1bp followed by the “rescue” experiment through microinjection of exogenous *Mad2l1bp* cRNAs. To this end, we first designed three specific “siRNAs” against *Mad2l1bp* in order to knock down the *Mad2l1bp* mRNA levels in oocytes. Although we have repeatedly validated that one of the three siRNAs was able to reduce the *Mad2l1bp* mRNA levels by 70% in human somatic cells *in vitro*, this siRNA failed to deplete *Mad2l1bp* mRNA significantly after injection into the oocytes (data not shown). Next, we adopted an alternative strategy by overexpression of *Mad2* mRNA inducing meiotic MI arrest in GV oocytes and monitored the meiotic progression during later culture in M16 medium (Wassmann et al., 2003). We thus transcribed the *Mad2* cRNAs *in vitro* and injected the *Mad2* cRNAs into oocytes at GV stage. As expected, almost all the oocytes from the GV stage were trapped at the MI stage with no polar body extrusion under extended culture in M16 medium (Figure 3E, upper panel). This is consistent with previous studies showing that Mad2 overexpression stimulated the meiotic spindle checkpoint, leading to MI arrest in the oocytes. Using this model, we next simultaneously co-injected the *Mad2* cRNAs combined with *Mad2l1bp* cRNAs (WT vs p.R285* mutant) generated after transcription from the linearized plasmids. Consistently, the majority of GV oocytes supplemented with the WT *Mad2l1bp* cRNAs developed beyond meiosis I, as judged by the PBE (Figure 3E and 3F). In contrast, the oocytes supplemented with the p.R285* *Mad2l1bp* cRNAs rarely extruded the polar body, suggesting that the oocyte maturation was impeded at the MI stage, and the MAD2L1BP variant lost its function *in vivo* as compared with its WT counterpart (Figure 3E and 3F).

**Figure 3.**
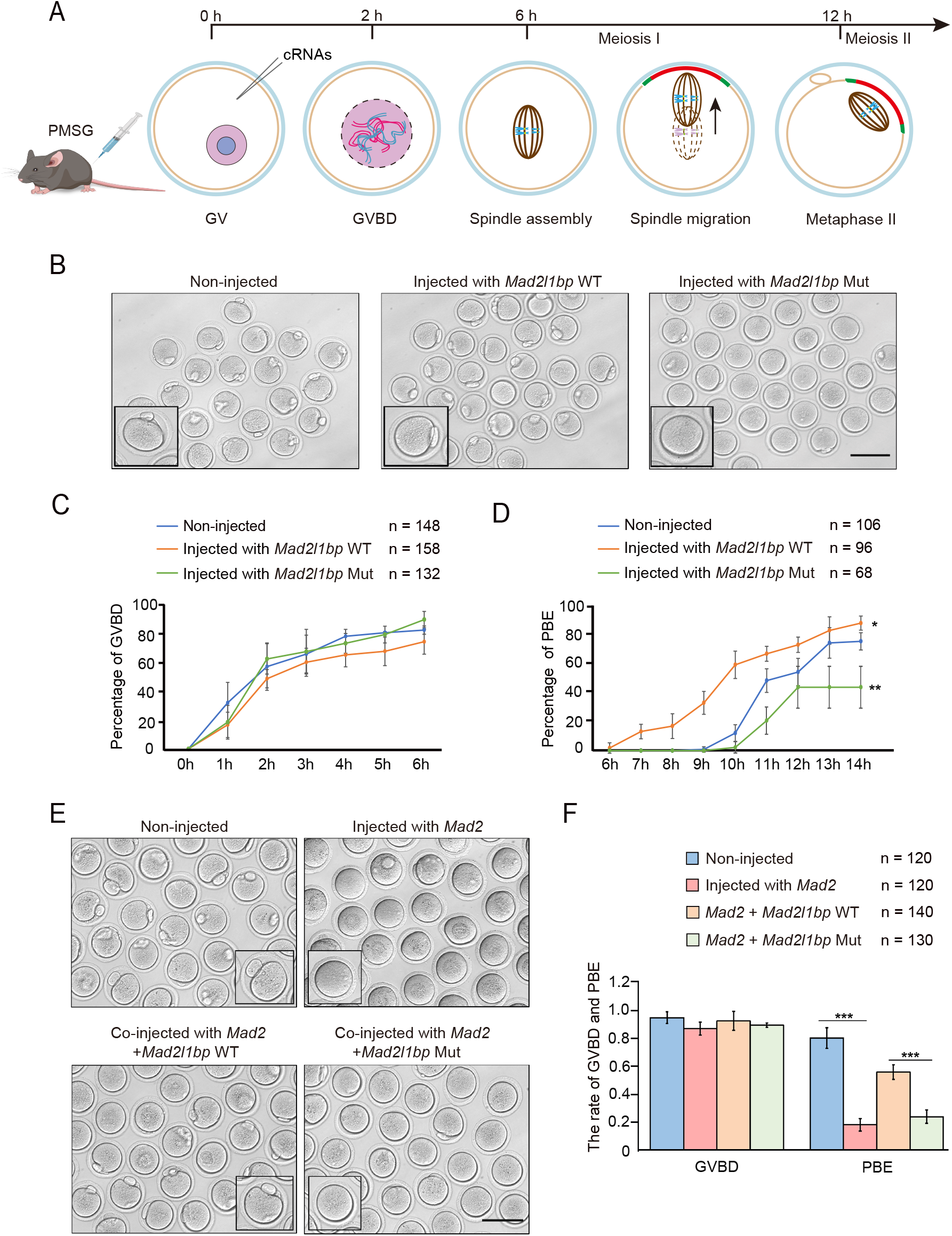
cRNA microinjection of *Mad2l1bp* accelerated or rescued the meiotic division in mouse GV oocytes. (A) A schematic diagram depicting the time points of distinctive events through meiotic progression in mouse oocytes. cRNAs were microinjected into GV oocytes after 44 hr of PMSG priming. (B) Representative morphology of oocytes after microinjection with cRNAs encoding mouse *Mad2l1bp WT* and *Mad2l1bp Mut* (equivalent to human p.R285*). Non-injected group was treated similarly to the other two groups except without cRNAs. Images were taken 14 hr following release from milrinone inhibitor. Scale bar, 100 μm. (C and D) Kinetic recordings showing the percentages of oocytes with GVBD (C), and the oocyte Polar Body Extrusion (PBE) rate (D) through the time-lapse imaging experiment. The total numbers of oocytes microinjected were labelled as indicated. The experiments were performed in biological triplicates. Data are presented as mean±SEM. (E) Representative morphology of oocytes after cRNA co-microinjection of Mad2 supplemented with WT or *Mad2l1bp* Mut (equivalent to human p.R285*) into the mouse GV oocytes; Images were taken 14 hr after release from milrinone. Scale bar, 100 μm. (F) Comparison of the percentages of oocytes with GVBD and PBE among the four microinjection groups in (E). The total numbers of oocytes microinjected were labelled as indicated. The experiments were performed in biological triplicates. The statistics were performed with *Student’s t test*. Data are presented as mean±SEM. “***” indicates p<0.001.

### Rescue of human oocyte meiotic arrest by microinjection of WT *MAD2L1BP* cRNA

The microinjection experiments performed in mouse oocytes as described above corroborated the *loss-of-function* effect of p.R285* *MAD2L1BP* variant. We next sought to determine whether we can recapitulate similar results in the frozen oocytes from the patients. We successfully recovered eight frozen oocytes with MI arrest from the first patient. Six oocytes were exploited for the microinjection of full-length *MAD2L1BP* cRNAs. Intriguingly, we attained four oocytes with PB1 after culture in G-MOPS medium for *in vitro* maturation, indicative of the successful completion of meiosis I in the patient’s oocytes when supplemented with exogenous intact *MAD2L1BP* cRNAs (Figure 4A). In contrast, the other two recovered oocytes with sham microinjection failed to extrude the polar body as a control (Figure 4A). This evidence manifests the p.R285* MAD2L1BP as a causative variant underlying oocyte MI arrest in the patient, and hints a possible treatment strategy by supplementation with exogenous *MAD2L1BP* cRNAs. Of note, the brother of the female patient from family 1 (Figure 1A) was also infertile. Examination of his semen sample showed that he had a markedly declined number of sperm, some of which exhibit aberrant head morphology, reminiscent of a possible role of MAD2L1BP in spermatogenesis as well (Figure 1-figure supplement 2, Supplementary file 1A).

**Figure 4.**
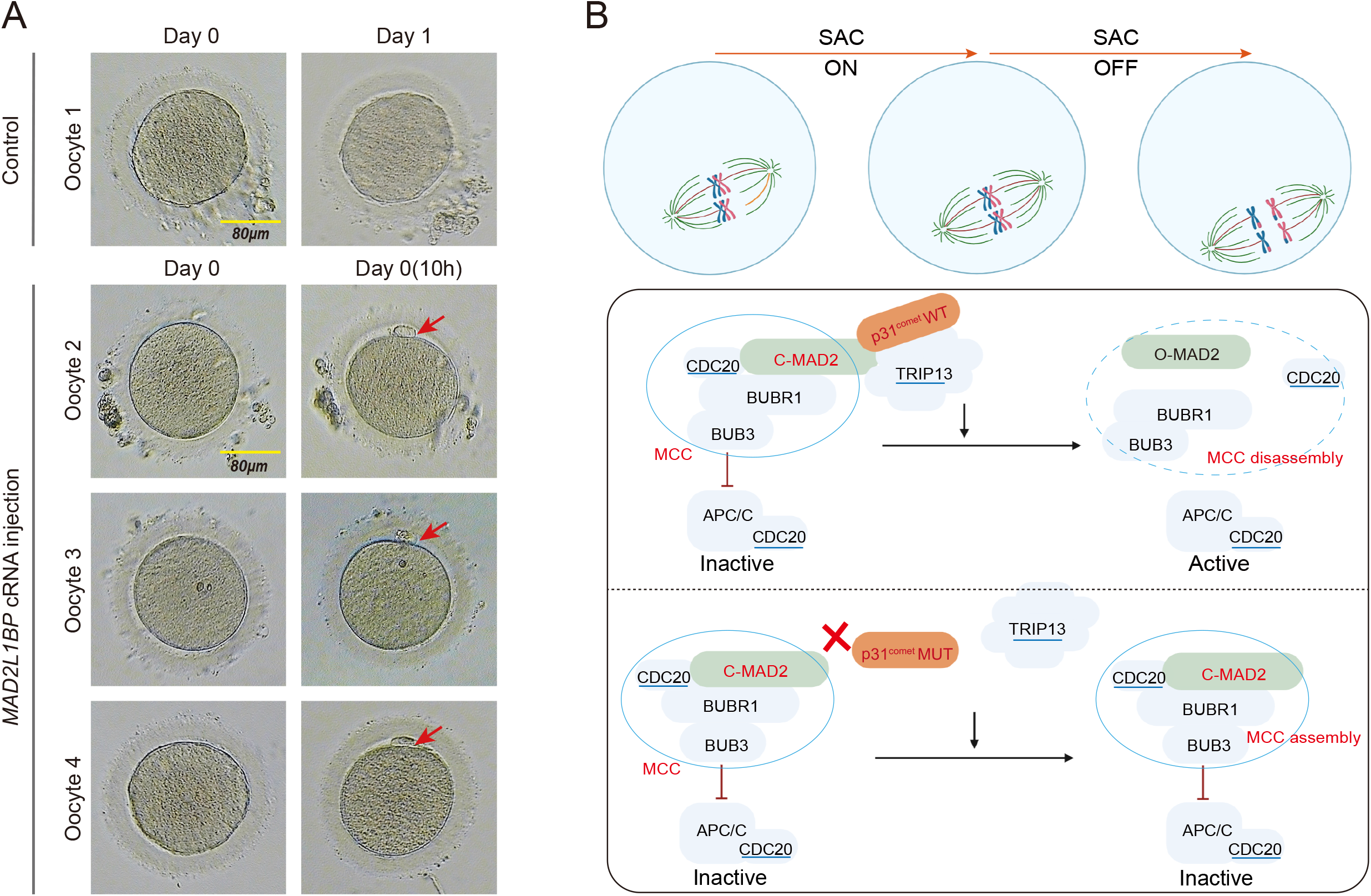
Meiotic Rescue of the MI-arrested frozen human oocytes by *MAD2L1BP* cRNA microinjection. (A) The resumption of polar body extrusion through *MAD2L1BP* cRNA microinjection into the frozen oocytes of the patient in Family 1 (II-1). The extrusion of the first polar body indicates the completion of meiosis I of the oocytes. In the control group, the retrieved MI oocytes remained arrested at MI after sham microinjection (upper panel). Six oocytes from the patient in Family 1(II-1) were microinjected with full-length *MAD2L1BP* cRNAs, of which four successfully finished extrusion of PB1 (lower panel). The red arrows point to PB1. Scale bar, 80 μm. (B) A summarized working model depicting how MAD2L1BP(p31^comet^) deficiency causes human oocyte meiotic arrest. The mitotic checkpoint complex (MCC) is well-known to comprise four core members: BUBR1, BUB3, CDC20 and MAD2. MAD2L1BP(p31^comet^) is known as the core adaptor that bridges the interaction between MAD2 and TRIP13(Alfieri et al., 2018; Yang et al., 2007). The MAD2L1BP (p31^comet^)-mediated interplay between MAD2 and TRIP13, known as the MAD2·MAD2L1BP(p31^comet^)·TRIP13 axis, disassembles the MCC signaling, which consequently drives the meiotic progression during oocyte meiosis. Top panel: The spindle assembly checkpoint (SAC) is switched “ON” until all the chromosome kinetochores are correctly connected to the spindles. Bottom panel: In the presence of WT MAD2L1BP, SAC is turned “OFF” through the recruitment of MAD2-MAD2L1BP-TRIP13 signaling to silence MCC, leading to the meiotic cell-cycle progression. By contrast, in the presence of mutated MAD2L1BP, MCC is unable to be silenced, resulting in persistent SAC activation and oocyte meiotic arrest.

## Discussion

To ensure the fidelity of chromosome segregation, the cells, including the oocytes, have evolved an exquisite surveillance machinery – the spindle assembly checkpoint (SAC), which represses the downstream anaphase-promoting complex (APC/C) activity until all chromosomes are correctly aligned and attached to the bipolar spindles through the kinetochores (Liu and Zhang, 2016) (Figure 4B). Central to the SAC is the physical presence of the mitotic checkpoint complex (MCC), which is comprised of four core components: BUBR1, MAD2, BUB3 and the APC activator CDC20. These players are phylogenetically conserved in metazoan species, and mouse models have provided unambiguous evidence showing their indispensable roles in cell cycle progression (Lara-Gonzalez et al., 2021; Liu and Zhang, 2016). In response to the kinetochores improperly attached to the spindles in the prometaphase, the SAC will be activated, which elicits a hierarchical recruitment of proteins to the outer kinetochores (Figure 4B). This drives the conversion of Mad2 from its inactive, open conformer (O-Mad2) to its closed, active form (C-Mad2) through a generally accepted “Mad1-Mad2 templating” mechanism for cytosolic propagation of spindle checkpoint signal(Lara-Gonzalez et al., 2021). Genetic mouse models have shown that Mad2 is the core player of SAC signaling – inactivation of Mad2 caused the accelerated mitotic exit whereas overexpression of Mad2 induced precocious activation of SAC in somatic cells and meiotic metaphase I arrest in oocytes, suggesting MAD2 is functional and required for normal oocyte meiotic progression(Lara-Gonzalez et al., 2021; Niault et al., 2007; Wassmann et al., 2003). Once the kinetochore-spindle attachment is satisfactorily achieved, the inhibitory signaling from the SAC must be timely quenched to allow anaphase progression. This is primarily achieved through MAD2 recognition by MAD2L1BP, which further recruits AAA+ ATPase, TRIP13, that binds and subsequently dissembles the MCC in an ATP-dependent manner (Alfieri et al., 2018; Brulotte et al., 2017). Conversely, deficient SAC disassembly, for example, *MAD2L1BP* mutations in this study, disrupts the MAD2·MAD2L1BP(p31^comet^)·TRIP13 axis, leading to sustained SAC activation and consequently the oocyte maturation arrest (Figure 4B, low panel).

Intriguingly, recent studies have identified pathogenic biallelic variants in *TRIP13* causing female primary infertility owing to the oocyte maturation arrest predominantly at MI stage(Zhang et al., 2020). Furthermore, a spectrum of deleterious variants in *CDC20*, the core component of MCC, have also been discovered, that account for human oocyte meiotic arrest and early embryonic failure (Zhao et al., 2020). To the best of our knowledge, this study is the first report that identified and characterized causative *loss-of-function* variants in *MAD2L1BP* underlying human oocyte maturation arrest. Consistent with the abnormalities in *MAD2L1BP-null* oocyte meiotic division, *Mad2l1bp* KO cells exhibited mitotic delay and lagging chromosomes (Choi et al., 2016). Moreover, in order to recapitulate the deleterious effect of the human *MAD2L1BP* variants in mice, we have attempted to generate the knockin (KI) mouse model that mimic the equivalent human mutations. However, *Mad2l1bp-*deficient mice died perinatally owing to the neonatal hypoglycemia resulting from the liver glycogen shortage, rendering us fail to get the homozygous KI offspring (Choi et al., 2016). Taken together, our study adds novel pathogenic variants of *MAD2L1BP* to the gene mutational spectrum underlying human oocyte maturation arrest, and likely provides novel therapeutic avenues for infertility treatment in the future.

## Materials and Methods

### Key resources table

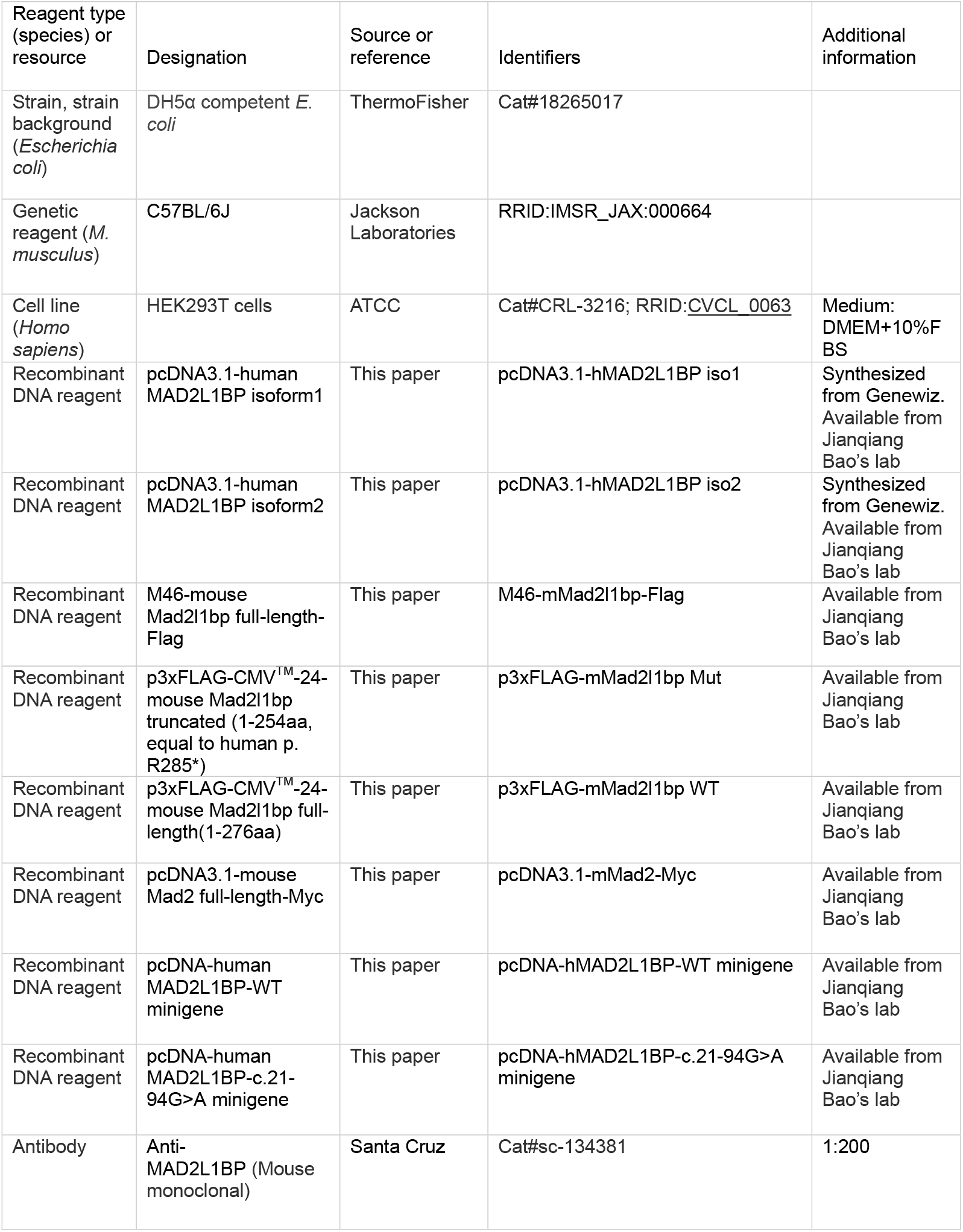
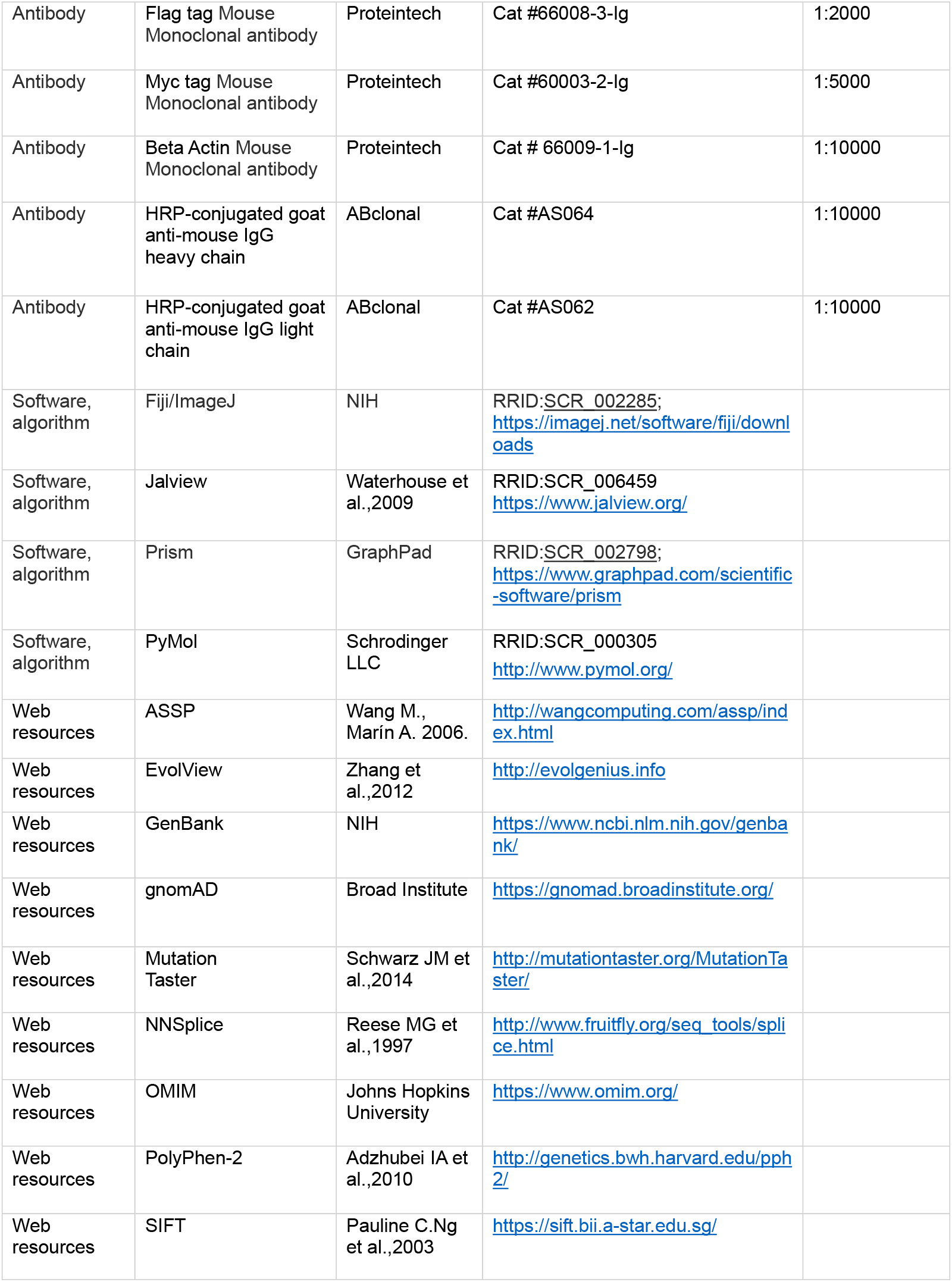

### Patients

A total of 50 primary infertile females undergoing recurrent IVF/ICSI failure due to complete oocyte maturation arrest were recruited between July 2014 and October 2021, from the First Affiliated Hospital of USTC (University of Science and Technology of China) and the Center for Reproductive Medicine, Shandong University. All of them had a normal karyotype (46, XX). Peripheral blood samples were obtained from all affected individuals and their available family members. Written informed consent was obtained from the participants. This study was approved by the biomedical research ethics committees of the First Affiliated Hospital of USTC (University of Science and Technology of China) and Shandong University.

### Whole-exome sequencing (WES) and data analysis

WES was performed using the DNA extracted from the periphery blood samples of the probands and the family members. Exonic DNA libraries were prepared using the Agilent Human SureSelect All Exon V6 kit and exome sequencing was performed on an Illumina NovaSeq 6000 platform. Clean sequencing reads were aligned to the human reference sequence (hg19). Sequence variants, including single-nucleotide variants (SNVs) and small insertions or deletions (indels), were annotated by the ANNOVAR pipeline. Common variants (defined as a minor allele frequency (MAF) above 1% in public databases: 1000 genome, dbSNP, ESP6500, gnomAD, or ExAC) were excluded. SNVs and indels were classified by position as intergenic, 5’ UTR, 3’UTR, intronic, splicing, or exonic. Exonic variants were then classified by predicted amino acid change as a stopgain, missense, synonymous, frameshift, indel or inframe, or possible splicing variants. For coding or possible splicing variants, the conservation at the variant site and the potential effect on protein function were evaluated with *in-silico* tools: SIFT, PolyPhen-2, MutationTaster, NNSplice, and ASSP (Table 3). Initially, we filtered genes that had predicted deleterious variants in at least two unrelated affected women and had not been previously reported (Table 2-source data 1).

### Sanger sequencing of the candidate variants

The DNA samples were extracted from the peripheral blood from the patients and their available family members. Sanger sequencing of all the coding regions, 200 bp of the flanking intron splicing sites and the promoter region (1223 bp upstream of the start codon) of *MAD2L1BP* were performed in the three probands and all their available family members. qPCR analysis of all the coding regions of *MAD2L1BP* was also performed to screen for small copy number variants (CNVs) in the patient (F2:II-1) and her family members. See primers in Supplementary file 1B.

### Plasmid construction

Total RNA was extracted from 10 ovaries from female mice between 4∼6 weeks of age using TRIzol following the manufacturer’s protocol. First-strand cDNA was synthesized using a ProtoScript II cDNA first strand kit (NEB) with 1 μg of total RNA. The full-length and truncated coding sequences of mouse *Mad2l1bp* were amplified and cloned into the p3xFLAG-CMV™-24 vector. cDNA fragment encoding mouse *Mad2* was PCR amplified and inserted into the pcDNA3.1 vector with a Myc tag. Human *MAD2L1BP* coding sequence were synthesized from Genewiz and cloned into the pcDNA3.1 vector. All construct sequences were verified by Sanger sequencing.

### Real-time quantitative PCR

Total RNA was extracted using a SPARKeasy Frozen whole blood total RNA Kit. 0.5 μg RNA from each sample was used for reverse transcription with a RevertAid First Strand cDNA Synthesis Kit (Thermo scientific). qPCR was performed with ChamQ Universal SYBR qPCR Master Mix (Vazyme) using a Roche LightCycler 96 machine. Relative mRNA levels were calculated by normalizing to the levels of internal GAPDH control. The qPCR primers are shown in Supplementary file 1B.

### Western blotting

Protein extracts were denatured by heating for 10 min at 95 °C in SDS–PAGE sample loading buffer. Proteins were separated by gel electrophoresis, followed by wet transfer to polyvinylidene difluoride (PVDF) membranes (Millipore). After blocking with 5% nonfat milk diluted in phosphate buffered saline supplemented with 0.05% Tween 20 for 1 hour, membranes were probed with primary antibodies against MAD2L1BP (1:200, Santa Cruz, Cat #sc-134381), Flag (1:2000, Proteintech, Cat #66008-3-Ig) and Myc (1:5000, Proteintech, Cat #60003-2-Ig). Beta-actin (1:10000, Proteintech, Cat # 66009-1-Ig) was used as internal control. The secondary antibodies were HRP-conjugated goat anti-mouse IgG heavy chain (1:10000, ABclonal, Cat #AS064) or HRP-conjugated goat anti-mouse IgG light chain (1:10000, ABclonal, Cat #AS062). Target proteins were detected using the ECL Western Blotting Detection Kit (Tanon) according to the manufacturer’s recommendation.

### Cell culture, plasmid transfection, and co-immunoprecipitation (CoIP)

293T cells were maintained in high-glucose DMEM (Gibco) medium supplemented with 10% fetal bovine serum (FBS; Gemini) and 1% penicillin-streptomycin solution (Gibco) at 37°C in a humidified 5% CO_2_ incubator. Transient transfections were performed using PEI reagent (Polyscience). Cells were collected 48 hours post transfection.

Immunoprecipitation assays were performed using 293T cells transfected with Flag-Mad2l1bp or Flag-Mad2l1bp(mut) and Myc-Mad2 plasmids as described above. Total protein lysates from transfected cells were prepared in lysis buffer [20 mM Tris-HCl (pH 7.5), 200 mM NaCl, 1% NP-40, and 1% protease inhibitor cocktail (APE-BIO)], and precleared with protein A Dynabeads (Invitrogen) and protein G Dynabeads (Invitrogen) for 1 hour at 4°C. The pre-cleared extracts (10% for input) were incubated with Flag antibody or mouse IgG overnight at 4°C. Protein A and protein G were added to the antibody-extracts mixture followed by incubation for one hour at 4°C. The beads were washed with lysis buffer for 4 times prior to elution with SDS sample buffer. Western blotting was conducted as described above.

### In vitro transcription and preparation of cRNAs for microinjections

To prepare cRNAs for human oocyte microinjection, constructs were linearized with the Not1 restriction enzyme, and the linearized products were purified by phenol (Tris-saturated):chloroform extraction and ethanol precipitation, and then dissolved in nuclease-free water. The purified DNA templates were transcribed *in vitro* using the T7 High Yield RNA Transcription Kit (Novoprotein, E131). Transcribed cRNAs were capped using the Cap1 Capping System (Novoprotein, M082) and poly(A) tails (∼100bp) were added using the E.coli Poly(A) Polymerase (Novoprotein, M012). cRNAs were purified by phenol(water-saturated):chloroform extraction and ethanol precipitated, and resuspended in nuclease-free water.

To prepare cRNAs for mouse oocyte microinjection, expression vectors were linearized and subjected to phenol (Tris-saturated):chloroform extraction and ethanol precipitation. The linearized DNAs were in vitro transcribed using the T7 message mMACHINE Kit (Invitrogen, AM1344). Transcribed mRNAs were added with poly(A) tails (∼200–250 bp) using the Poly(A) Tailing Kit (Invitrogen, AM1350), recovered by lithium chloride precipitation, and resuspended in nuclease-free water.

### Human oocyte collection and microinjection

Human oocytes (from control and affected individuals) were donated following informed consent. Oocytes from Family 1(II-1) were recovered and randomly divided into the control and cRNA-injected groups. About 5 pl *MAD2L1BP* cRNA solution (1000 ng/μl) was microinjected into the oocytes. After injection, oocytes were cultured for *in vitro* maturation and considered to be matured when PB1 was extruded.

### Mouse oocyte culture and microinjection

Animal care and experimental procedures were conducted in accordance with the Animal Research Committee guidelines of Zhejiang University (approval # ZJU20210252 to H.Y.F). The 21–23-day-old female mice were injected with 5 IU of PMSG and euthanized after 44 h. Oocytes at the GV stage were harvested in M2 medium (Sigma-Aldrich, M7167) and cultured in mini-drops of M16 medium (Sigma-Aldrich, M7292) covered with mineral oil (Sigma-Aldrich, M5310) at 37°C in a 5% CO_2_ atmosphere.

For microinjection, fully grown GV oocytes were incubated in M2 medium with 2 μM milrinone to inhibit spontaneous GVBD. All injections were performed using an Eppendorf transferman NK2 micromanipulator. Denuded oocytes were injected with 5–10 pl samples per oocyte. The concentration of all injected RNAs was adjusted to 1200 ng/μl. After injection, oocytes were washed and cultured in M16 medium at 37°C with 5% CO_2_.

### Minigene assay

Minigene analysis was performed in 293T cells. Amplicons spanning exons 1–3 along with ∼200 bp of the flanking intronic sequences were PCR-amplified using the genomic DNA from the normal control and were inserted into the pcDNA vector. Site-directed mutagenesis was performed to introduce the mutation c.21-94G>A. The minigene constructs (c.21-94G>A or WT) were transfected into cultured 293T cells using PEI reagent (Polyscience). Cells were harvested 48 h after transfection and total RNA was extracted using TRIzol following the manufacturer’s protocol. RT-PCR was performed using the indicated primers to amplify the target region (See Supplementary file 1B for primers sequences).

### Molecular modeling and evolutionary conservation analysis

MAD2L1BP mutations were mapped using PyMOL software according to the crystal structure of the Mad2/p31(comet)/Mad2-binding peptide ternary complex (PDB ID: 2QYF). Evolutionary conservation analysis was performed with Jalview software, and processed on the Evolview website.

### Evaluation of oocyte phenotypes and sperm phenotypes

The stage of oocyte maturation was assessed under a light microscope as described previously(Huang et al., 2018). Semen analysis and papanicolaou staining were performed according to the protocol of the World Health Organization 2010 guidelines.

### Analysis of the expression pattern of MAD2L1BP and MAD2 in human follicles and mouse oocytes

RNA-seq data was downloaded from GSE71434(Zhang et al., 2016) and GSE107746 (Zhang et al., 2018). Reads were mapped to the mouse genome (mm10) or human genome (hg38) using STAR. Relative expression quantification of genes and transcripts was performed using RSEM.

## Statistical analysis

Data are presented as the mean ± standard error of the mean (SEM). Most experiments were repeated at least three times. The results for two experimental groups were compared by two-tailed unpaired Student’s t-tests. Statistically significant values of P < 0.05, P < 0.01, and P < 0.001 by two-tailed Student’s t-test are indicated by asterisks (*), (**), and (***), respectively. “n.s.” indicates nonsignificant.

## Figure supplement legends

**Figure 1- figure supplement 1.**
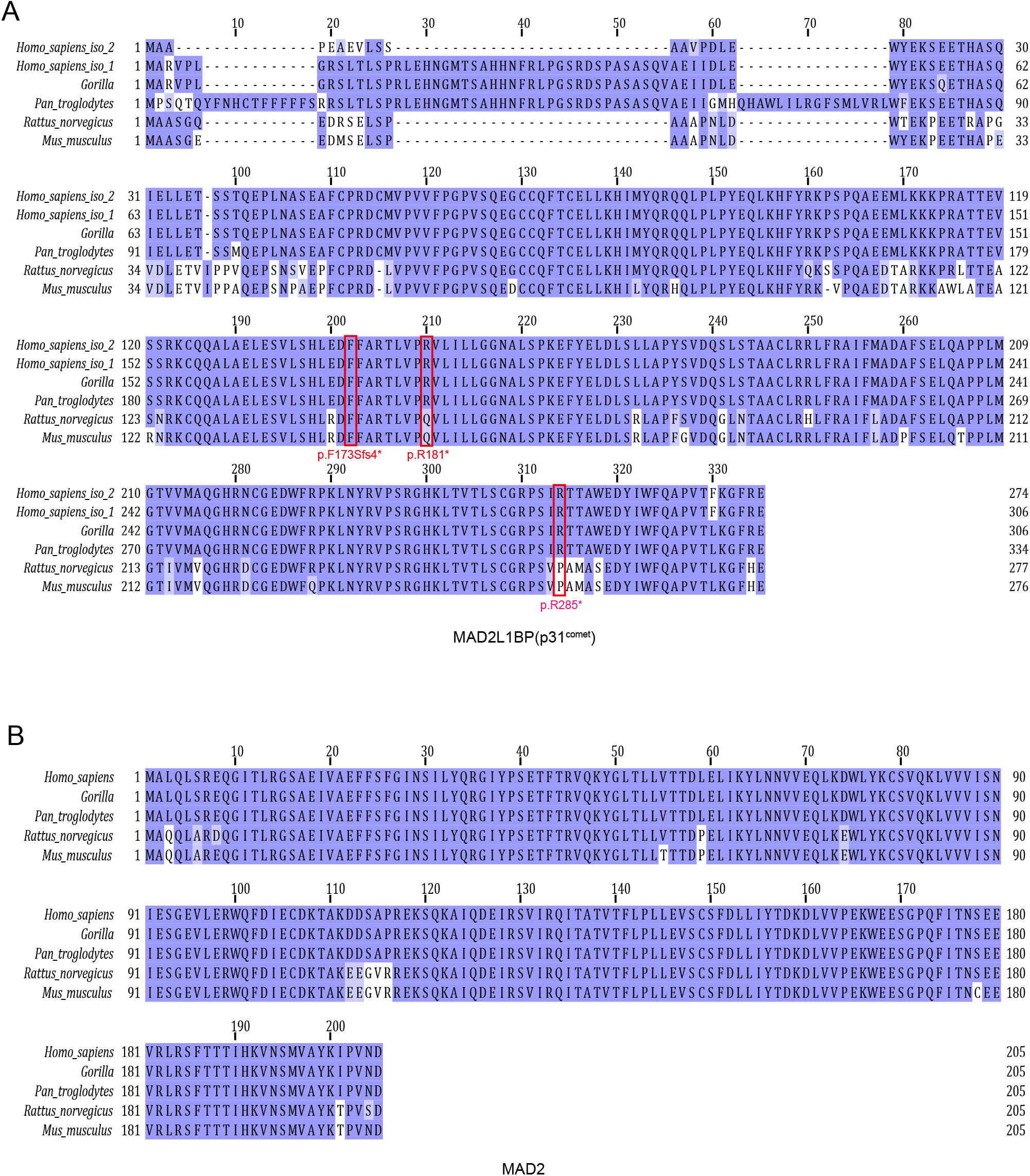
Multi-sequence Alignment of Amino Acids for MAD2L1BP (p31^comet^) (A) and its interacting partner MAD2 (B) orthologs from Eutherian Species with Jalview. Pathogenic mutations in affected individuals reported in this study are marked in red box.

**Figure 1- figure supplement 2.**
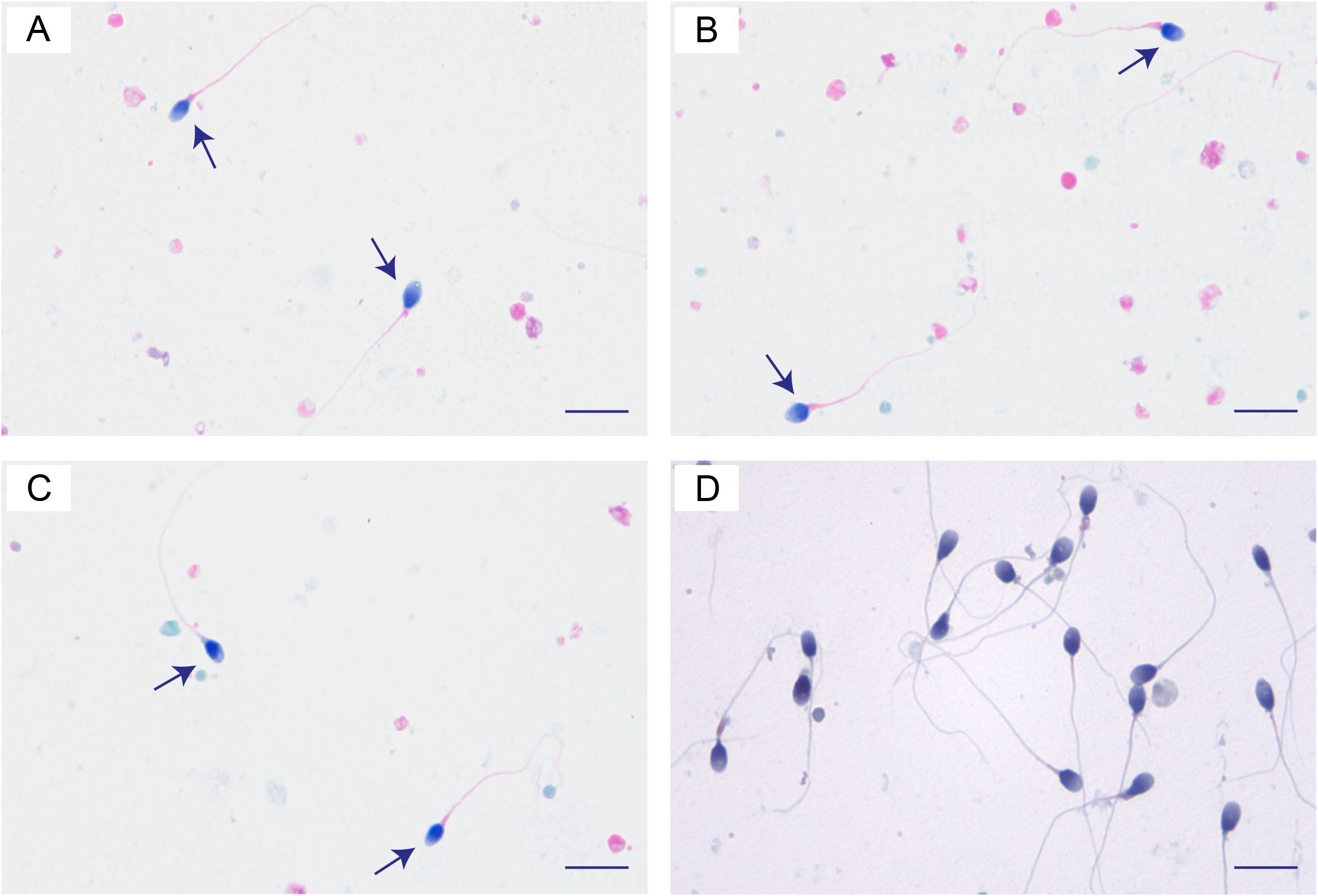
Semen Analysis of the Individual II-2 in Family 1 with a Homozygous Mutation (c.853C>T [p.R285*]) in *MAD2L1BP*. (A-C) Photomicrographs of the ejaculated semen smear stained by Papanicolaou. Few normally-shaped spermatozoa can be observed in the ejaculated semen from the individual II-2 in Family1. (D) The control sperm morphology by papanicolaou staining from a fertile man. Scale bars, 10 μm. See Table S1 for semen parameters of the individual II-2 in Family1.

**Figure 2- figure supplement 1.**
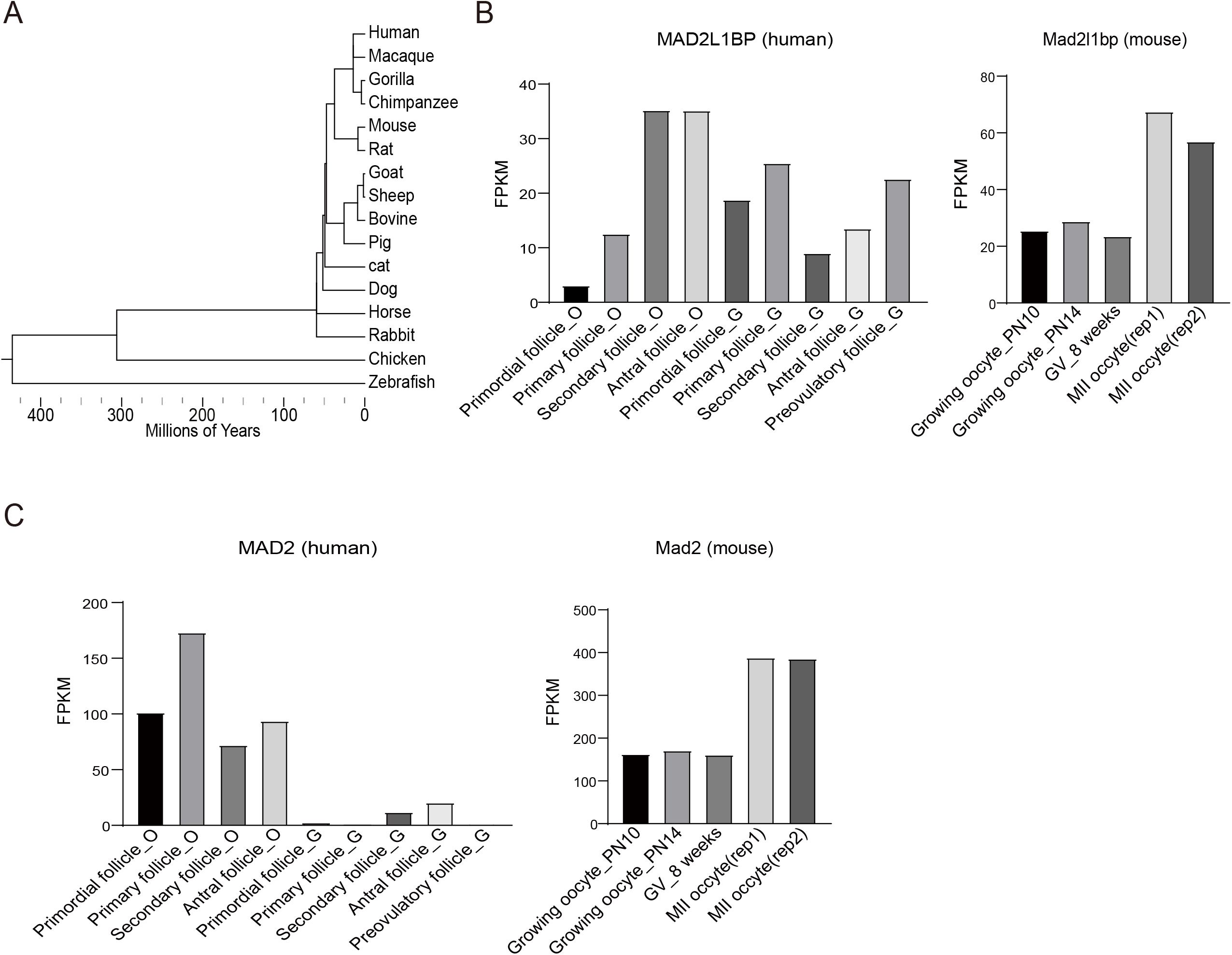
Phylogenetic Tree of *MAD2L1BP* and mRNA Expression Pattern of *MAD2L1BP* and *MAD2* in Human Follicles and Mouse Oocytes. (A) Evolution of the *MAD2L1BP* gene orthologs in metazoan species. Sequence conservation analysis was performed with Jalview software, and processed on Evolview website. (B-C) mRNA expression levels of Mad2l1bp (B) and Mad2l1(Mad2) (C) in developing human follicles and mouse oocytes. RNA-seq data was downloaded from GSE71434 and GSE107746. Reads were mapped to the mouse genome (mm10) or human genome (hg38) using STAR. Relative expression quantification of genes and transcripts was performed using RSEM. PN, postnatal; GV, germinal vesicle; MII, metaphase II; rep, replicate. O, oocyte; G, Granulosa cell.

**Figure 2- figure supplement 2.**
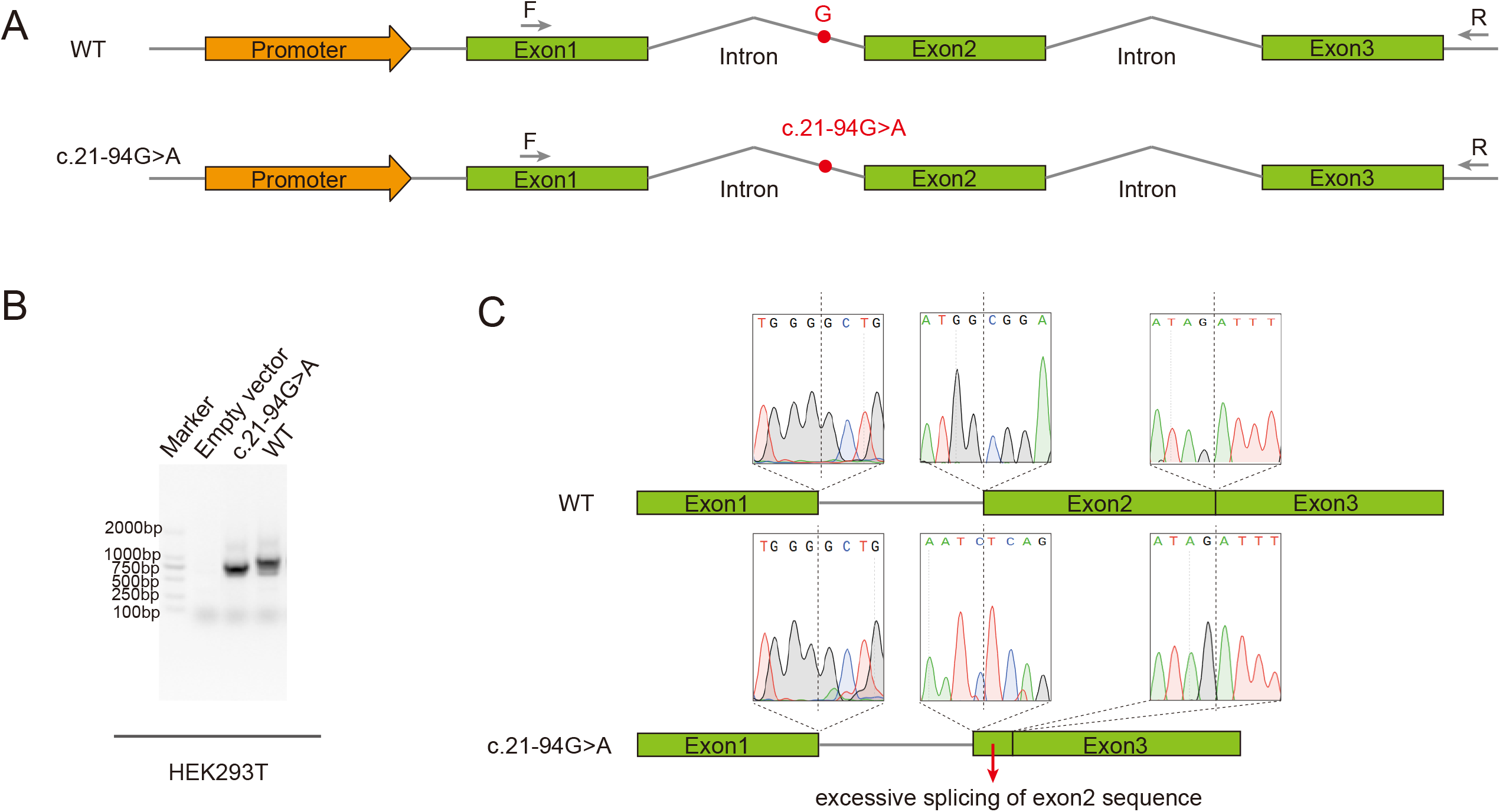
Minigene assay showing the aberrant alternative splicing of *Mad2L1BP* exons associated with the variant c.21-94G>A from the family 2. (A) Schematic diagram of minigene splicing assay. The G→A substitution in intron 1 produced a novel acceptor site for Exon 2, leading to the generation of a premature termination codon (PTC). HEK293T cells were transfected with minigene constructs (WT or c.21-94G>A), RT-PCR was performed with the primers as indicated. (B) Different splicing patterns in WT and c.21-94G>A group were shown by the agarose gel. (C) Representative Sanger sequencing chromatogram validating the excessively splicing of the exon 2 in *MAD2L1BP* mRNA with c.21-94G>A and thus caused deletion of large fragment of MAD2L1BP protein.

## Source data legends

**Figure 2-source data 1:**

Blots for detecting Myc-MAD2, Flag-MAD2L1BP WT, Flag-MAD2L1BP MUT protein expression.

**Figure 2-source data 2:**

Co-IP blots for detecting interaction between MAD2 and MAD2L1BP WT or MAD2L1BP MUT

**Figure 3-source data 1:**

cRNA microinjection of full-length or truncated *Mad2l1bp* uncovered their discordant roles in driving the extrusion of polar body 1 (PB1) in mouse oocytes.

**Figure 3-source data 2:**

Co-injection of *Mad2* cRNAs with *Mad2l1bp* cRNAs (WT vs p.R285* mutant) indicated the Mad2l1bp variant lost its function in vivo as compared with its WT counterpart.

**Table 2 -source data 1:**

Homozygous and compound heterozygous variants identified by WES that survived filtering in patient F1: II-1, F2: II-1 and F3: II-1.

## Data availability

All data generated or analyzed during this study are included in the manuscript and supporting files. Source Data files have been provided for Figure 2, Figure 3 and Table 2.

## Supplementary File

The Supplementary file include Supplementary file 1A and 1B.

## Declaration of interests

The authors declare no competing interests.

## Materials availability statement

Plasmids constructed in this paper are available from Jianqiang Bao’s lab upon request.

## Acknowledgments

This study was supported by the grants from National Natural Science Foundation of China (81801440, 82192874, 31871509, 31970793, 32170856), the Ministry of Science and Technology of China (2019YFA0802600), the Fundamental Research Funds for the Central Universities” (WK2070000156, WK9100000032) and Startup funding (KY9100000001). The authors thank Prof. Lei Wang (Fudan University) for discussion with this manuscript. We thank Dr.Jianming Zeng(University of Macau), and all the members of his bioinformatics team, biotrainee, for generously sharing their experience and codes. The Use of the biorstudio high performance computing cluster(https://biorstudio.cloud) at Biotrainee and The shanghai HS Biotech Co.,Ltd for conducting the research reported in this paper.

